# Feedback Determines the Structure of Correlated Variability in Primary Visual Cortex

**DOI:** 10.1101/086256

**Authors:** Adrian G. Bondy, Ralf Haefner, Bruce G. Cumming

## Abstract

The variable responses of sensory neurons tend to be weakly correlated (spike-count correlation, *r*_*sc*_). This is widely thought to reflect noise in shared afferents, in which case *r_sc_* can limit the reliability of sensory coding. However, it could also be due to feedback from higher-order brain regions. Currently, the relative contribution of these sources is unknown. We addressed this by recording from populations of V1 neurons in macaques performing different discrimination tasks involving the same visual input. We found that the structure of *r*_*sc*_(the way *r*_*sc*_ varied with neuronal stimulus preference) changed systematically with task instruction. Therefore, even at the earliest stage in the cortical visual hierarchy, *r*_*sc*_ structure during task performance primarily reflects feedback dynamics. Consequently, previous proposals for how *r*_*sc*_ constrains sensory processing need not apply. Furthermore, we show that correlations between the activity of single neurons and choice depend on feedback engaged by the task.

---

Judgments made about sensory events (i.e. perceptual decisions) rely on the spiking discharge of sensory neurons. For this reason, there has been longstanding interest in the observation that this discharge tends to be variable given a fixed stimulus^1,2^. In principle, this variability could confound perceptual judgments, impairing the fidelity of sensory information in the brain. Even worse, this variability tends to be weakly correlated amonst sensory neurons (spike-count correlation; *r*_*sc*_)^3^, meaning it cannot trivially be averaged away^4^. For this reason, *r*_*sc*_ is widely referred to as “correlated noise”^5–8^.

This way of thinking has underlied several influential lines of research in systems neuroscience. One has sought to understand the magnitude of the perceptual impairment introduced by *r*_*sc*_ in different behavioral contexts^5,8–15^. When *r*_*sc*_ is distributed in such a way that correlated fluctuations mimic the sensory events being detected or discriminated, it could severely impair perceptual accuracy^11,15,16^. A related line of research has sought to understand how correlated variability affects the choices subjects make in perceptual discrimination tasks from trial to trial^17–19^. These studies have shown that *r*_*sc*_ structure can give rise to a weak correlation between variability in single neurons and perceptual reports (Choice Probability; CP), consistent with the notion that CP observed in real neurons reflects the causal influence of correlated sensory neuronal variability on perception.

However, we currently know very little about the origin of *r*_*sc*_, making it unclear to what degree these conclusions are correct. A frequent (although typically unstated) assumption is that *r*_*sc*_ in sensory neurons is generated by shared variability in common afferent inputs. Consistent with this idea, *r*_*sc*_ correlates with the physical proximity and similarity in stimulus preference of neuronal pairs^8,20–23^, which are also predictive of the degree of feedforward input convergence. If this explanation is correct, it supports the traditional view of *r*_*sc*_ as “confounding noise” since it arises from stochastic processes in the sensory encoding pathway. However, the bulk of synaptic inputs to sensory cortical neurons are not strictly “feedforward” in nature^24,25^. Consequently, variation over time in shared inputs from downstream areas (i.e. “top-down”; “feedback”), may make a significant contribution to *r*_*sc*_. These signals may reflect endogenous processes like attention, arousal, or perceptual state, and could be under voluntary control. In principle, this source of correlated variability need not confound perceptual judgments, but instead reflect ongoing neuronal computations.

Several recent studies have shown that *r*_*sc*_ does change to some degree with task context^12,14,26,27^, suggesting a top-down component. These studies have shown that *r*_*sc*_ in populations of sensory neurons can either increase or decrease depending on attentional state or other task demands. However, prior studies have made only limited measures of *r*_*sc*_ structure and how this changes with task, yet these are critical for understanding how *r*_*sc*_ arises and how it relates to task performance. Furthermore, the relative magnitude of feedforward versus top-down contributions to *r*_*sc*_ has not been determined. It also unknown whether task-dependent changes in *r*_*sc*_ reflect an adaptive reduction of sensory noise or whether *r*_*sc*_ is, in the first instance, generated by variability over time in top-down inputs reflecting downstream computations.

In the present study, we used large-scale neuronal population recordings in behaving macaques, along with careful behavioral control and a novel analytical approach, to significantly advance our understanding of these fundamental questions. Subjects performed different orientation discrimination tasks using the same set of stimuli. The only difference between tasks was the set of orientations being discriminated. If *r*_*sc*_ primarily reflects noisy sensory encoding, it should be invariant to changes in the task given fixed retinal input. Alternatively, if it changes dynamically with the task, this would indicate that it reflects top-down signals. This experimental approach, inspired by a previous study^27^, was combined with large-scale population recordings, allowing us to estimate the full *r*_*sc*_ matrix – that is, how *r*_*sc*_ varies as a function of all possible combinations of pairwise orientation preference. This made it possible to directly infer which components were fixed and which changed with the task. Strikingly, we could not identify a component that remained fixed. Instead we observed a pattern of task-dependent changes that was highly systematic, and could be modeled as the effect of a single modulatory input that targets the two task-relevant subpopulations of V1 neurons in an alternating fashion across trials.

These data give unprecedented insight into the functional role of *r*_*sc*_ structure in task performance. First, they show that the task-dependent changes in *r*_*sc*_ structure appear to degrade the task performance of an ideal observer of V1 activity alone, because they mimic task-relevant stimulus changes. However, our discovery of the feedback origin of these correlations means that they need not degrade performance, and points to the possibility that they may instead be a signature of ongoing neuronal computations. Indeed, recent circuit models of perceptual inference predict feedback signals whose statistics reflect the subject’s prior beliefs about the task, yielding predictions which closely match our obervations^28,29^. Second, we show quantitatively that these feedback dynamics are the primary source of the choice-related activity we observed in V1, clarifying an ongoing debate^30^ about the interpretation of choice-related signals in sensory neurons. We conclude that *r*_*sc*_ in sensory neurons reveals less than previously thought about the encoding of sensory information in the brain, but potentially much more about the interareal computations underlying sensory processing.

## Results

We trained two rhesus monkeys (*Macaca mulatta*) to perform a two-alternative forced choice (2AFC) coarse orientation discrimination task (Fig. 1), used previously^31^. On a given trial, the subject was shown a dynamic, 2D filtered noise stimulus for 2 seconds, after which it reported the stimulus orientation by making a saccade to one of two choice targets (oriented Gabor patches). Different task contexts were defined by the pair of discriminandum orientations. The stimuli were bandpass filtered in the Fourier domain to include only orientations within a predetermined range. The stimulus filter was centered on one of the two task orientations and its orientation bandwidth was used to control task difficulty. We included 0%-signal trials, for which the stimuli were unfiltered for orientation (and thus the same regardless of context), to examine the effect of task context on *r*_*sc*_ in the presence of a fixed retinal input.

**Fig. 1.**
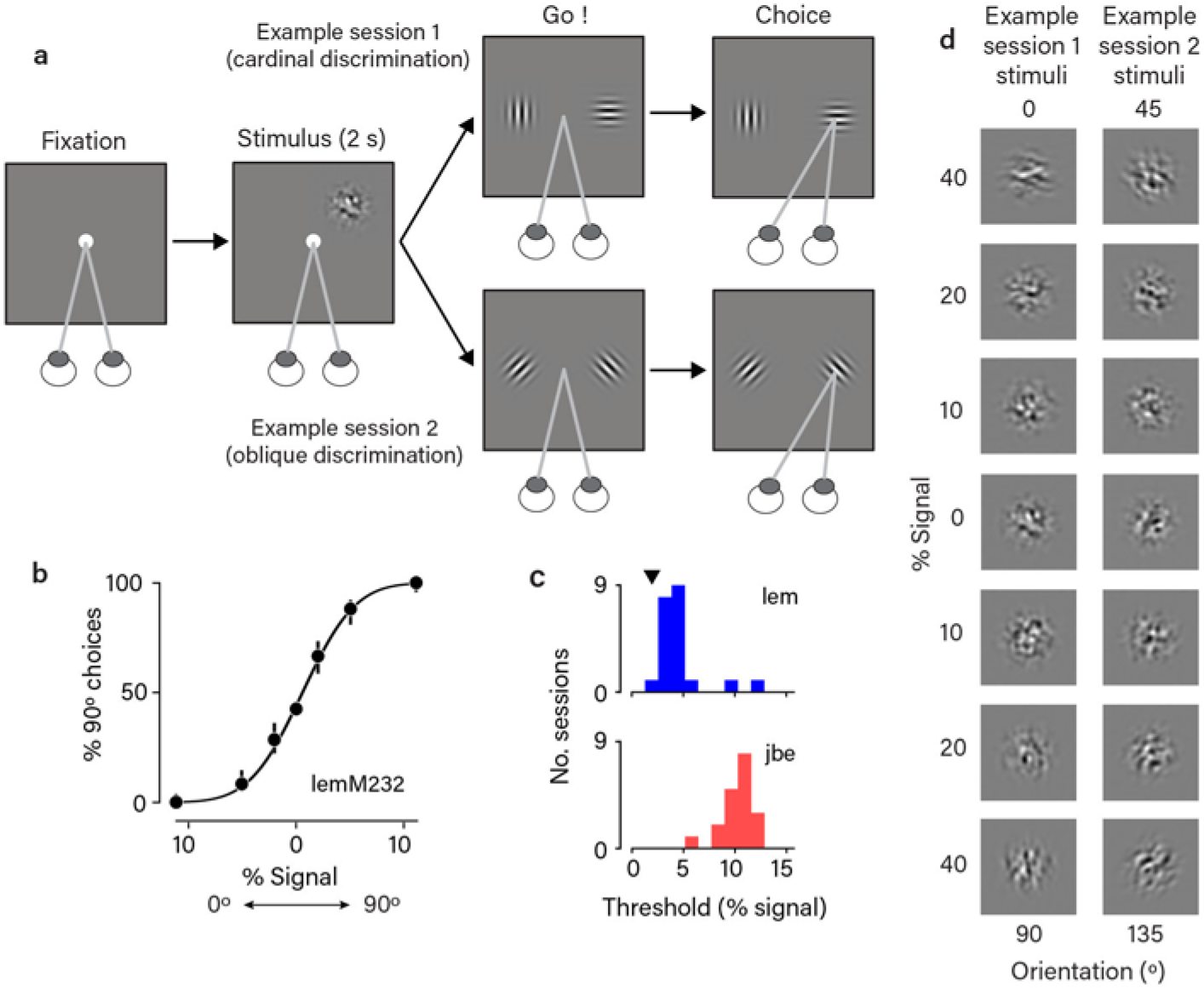
Orthogonal orientation discrimination task. **a.** Schematic 381 illustration of the task. Two example task contexts shown (cardinal and oblique discriminations). The task context was fixed in a given recording session, but varied across sessions. **b**. Psychometric function for monkey *‘lem’*, example session, n=1,354 trials. Black curve is a probit fit, and error bars are 95% confidence intervals around the mean (black points). **c**. Histograms showing the distribution of psychometric thresholds across sessions for the two subjects. Threshold is defined as the signal level eliciting 75% correct performance. Black triangle indicates the threshold corresponding to the example session in (b). **d**. Example single stimulus frames corresponding to the two example task contexts in (a). The stimuli consisted of dynamic, white noise filtered in the Fourier domain for orientation (see Methods). The filter was centered on one of the two task orientations and its bandwidth determined signal strength.

In order to detect any effect of task context on *r*_*sc*_ structure, it is critical that subjects based their choices on the presence of the correct orientation signals. To ensure this, we used psychophysical reverse correlation^31–33^ to directly measure the influence of different stimulus orientations on the subject’s choices (the “psychophysical kernel”). We found that subjects required multiple days of retraining after a change in the task context to fully update their psychophysical kernel. For this reason, we kept the task context fixed for the duration of each recording session, and only undertook recordings in a new task context after subjects had updated their kernel (Supplementary Fig. 1). This is a significant advance over past studies of the effect of task context on neuronal responses, which typically have not quantified the extent to which behavioral strategy truly matches task instruction.

We recorded spiking activity in populations of single V1 neurons using multi-electrode arrays while the subjects performed the task. We determined the preferred orientation of each neuron by measuring its response to oriented stimuli (see Methods) in separate blocks of trials during which subjects passively fixated. Neurons were excluded from analysis if they were not well orientation tuned. The final dataset includes 811 simultaneously recorded pairs from 200 unique cells across 41 recording sessions. For each pair, we calculated its *r*_*sc*_ value as the Pearson correlation between the set of trial-duration spike-counts across trials of the same stimulus condition. While measuring *r*_*sc*_ only across 0%-signal trials isolated any changes due to the task context, we found similar results within each signal level (Fig. 6). Therefore, to increase statistical power, we report *r*_*sc*_ values measured across all trials, after normalizing spike counts to remove the effect of stimulus drive on firing rates.

### *R_sc_* structure changes systematically with task context

Recording large populations gave us the power to measure the full “*r*_*sc*_ matrix”: that is, how *r*_*sc*_ varied as a function of all possible combinations of orientation preference. This is the first time that such detailed measures of *r*_*sc*_ structure have been made while animals perform a discrimination task. To assess the presence of task-dependent *r*_*sc*_ structure in the data, we we first divided the recording sessions into two groups based on the task context used (Fig. 2b). We estimated the smoothed, average *r*_*sc*_ matrix associated with each subset (Fig. 2a,c) by pooling *r*_*sc*_ values measured across the subset of sessions along with measures of the neuronal preferred orientation. Across both subsets of sessions, we observed a tendency towards higher values of *r*_*sc*_ for pairs of neurons with more similar orientation preferences (i.e. higher values closer to the diagonal of the matrix), consistent with numerous prior observations^3^ (Fig. 2d). Traditionally, such observations were presumed to reflect “limited-range correlations” that depend only on similarity in stimulus preference^5,9,10^, equivalent to a rotationally-symmetric (Toeplitz) correlation matrix. In contrast, in our data this was due to distinct patterns in the two matrices: we observed the highest values of *r*_*sc*_ amongst pairs that shared a preferred orientation close to a discriminandum, and the lowest values of *r*_*sc*_ tended to occur amongst pairs preferring opposite task orientations. Because the task context differed between the two subsets, this yielded matrices with a lattice-like pattern offset along the diagonal by an amount reflecting the task context. In other words, *r*_*sc*_ structure changed dramatically with task context, consistent with the presence of task-dependent feedback and inconsistent with a fixed *r*_*sc*_ structure primarily driven by sensory afferent noise.

**Fig. 2.**
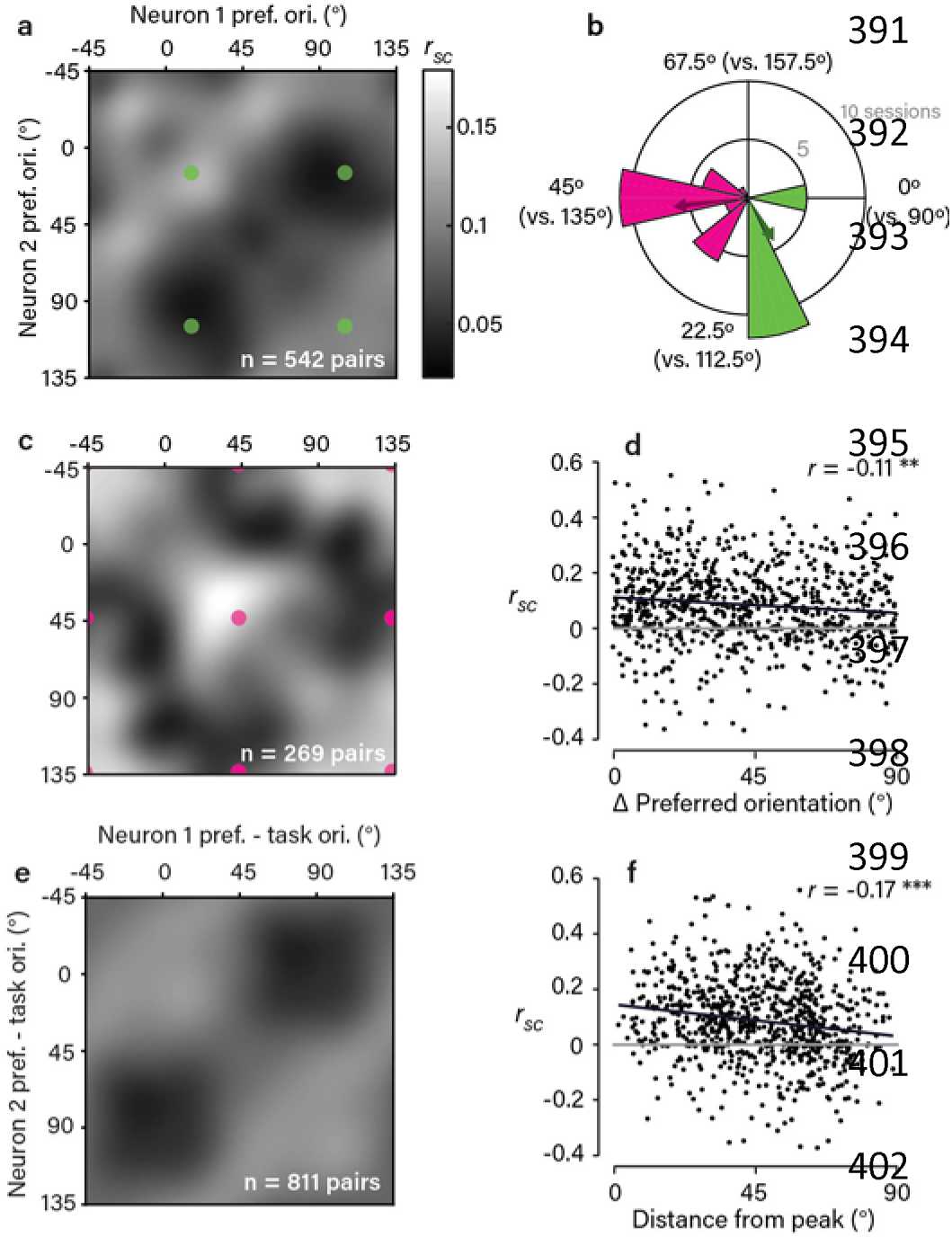
*R_sc_* structure in V1 depends systematically on task context. **a,c.**sObserved *r*_*sc*_ matrices for the two subsets of sessions grouped by task context, as indicated in (**b**). The matrices were obtained by pooling the set of *r*_*sc*_ measurements made within each subset and applying a von Mises smoothing kernel (approximating a 2D wrapped Gaussian with 15º s.d.). Colored dots correspond to pairs preferring the same or opposing task orientations. **b**. Polar histogram shows the distribution of task contexts used across sessions, with color indicating the division into two subsets. Note that the period is 90º because of the orthogonality of the discriminanda. Colored arrows indicate the mean task context associated with each subset. **d.** Scatter plot showing a weak, but significant, dependence of *r*_*sc*_ on the difference in preferred orientation of neuronal pairs (p=9*10^−4^, bootstrap test, one-sided). Black line is (type II) regression line and grey line corresponds to *r*_*sc*_=0. **e.** Average *r*_*sc*_ matrix observed across all session, shown in a task-aligned coordinate frame. Each pair’s preferred orientations are expressed relative to the task orientations (defined as 0º and 90º). Color scale as in (a). **f.** Scatter plot showing a significant dependence of *r*_*sc*_ on distance from the peak (0º/0º or 90º/90º) in the matrix in (**e**). This dependence was stronger than the dependence on difference in preferred orientation (r=-0.17, p=1.63*10^−6^, bootstrap test, one-sided), suggesting the task-aligned pattern we observed captures a more important feature of *r*_*sc*_ structure. Black and grey lines as in (d).

To summarize this task-dependent structure across the entire dataset (Fig. 2e) we expressed each neuron’s preferred orientation relative to the task orientations on its respective recording session, such that 0º and 90º always indexed the task orientations. This combined matrix clearly illustrates the task-dependent pattern of *r*_*sc*_ structure in the V1 population, a pattern that was consistent across both subjects (Supplementary Fig. 2). As in previous studies, there was a great deal of variability between individual *r*_*sc*_ values, even amongst pairs with similar orientation preferences and task (Fig. 2d,f) demonstrating that factors not considered here also contribute to *r*_*sc*_.

Importantly, we observed a different result during separate blocks of trials in the same recording sessions, during which the subject fixated passively for reward but the same set of stimuli was shown. During these blocks, the highest values of *r*_*sc*_ tended to occur along the diagonal, independent of orientation preference or task (Supplementary Fig. 3). This demonstrates that the task-dependent pattern observed during task performance depends on active task engagement, and cannot be explained, for instance, simply as an effect of adaptation to task experience. We performed a number of additional analyses to rule out any possibility that our findings could be explained merely as an effect of changing retinal input across task contexts, such as effects related to stimulus history or eye movements (see Supplementary Figs. 4-7). Taken together, these controls strengthen our interpretation that centrally-generated signals reflecting task engagement underlie the task-dependent *r*_*sc*_ structure we observed.

### Segregating fixed and task-dependent components of *r*_*sc*_ structure

Our dataset of *r*_*sc*_ measurements made in large, heterogeneous populations across diverse task contexts allowed us to directly estimate the *r*_*sc*_ structure that was fixed versus dynamically changing with task. To do this, we modeled the raw *r*_*sc*_ values using two structured components: 1) a fixed *r*_*sc*_ matrix describing the dependence of *r*_*sc*_ on pairwise orientation preference regardless of task, and 2) a task-dependent *r*_*sc*_ matrix capturing the dependence of *r_sc_* on pairwise orientation preference *relative* to the task orientations. We used ridge regression to find the form of these two component matrices that best predicted the raw *r*_*sc*_ measurements. To reduce the number of regressors without constraining the form these two components could take, we parametrized the matrices as 8x8 grids of basis functions (see schematic in Fig. 3a and Methods).

**Fig. 3.**
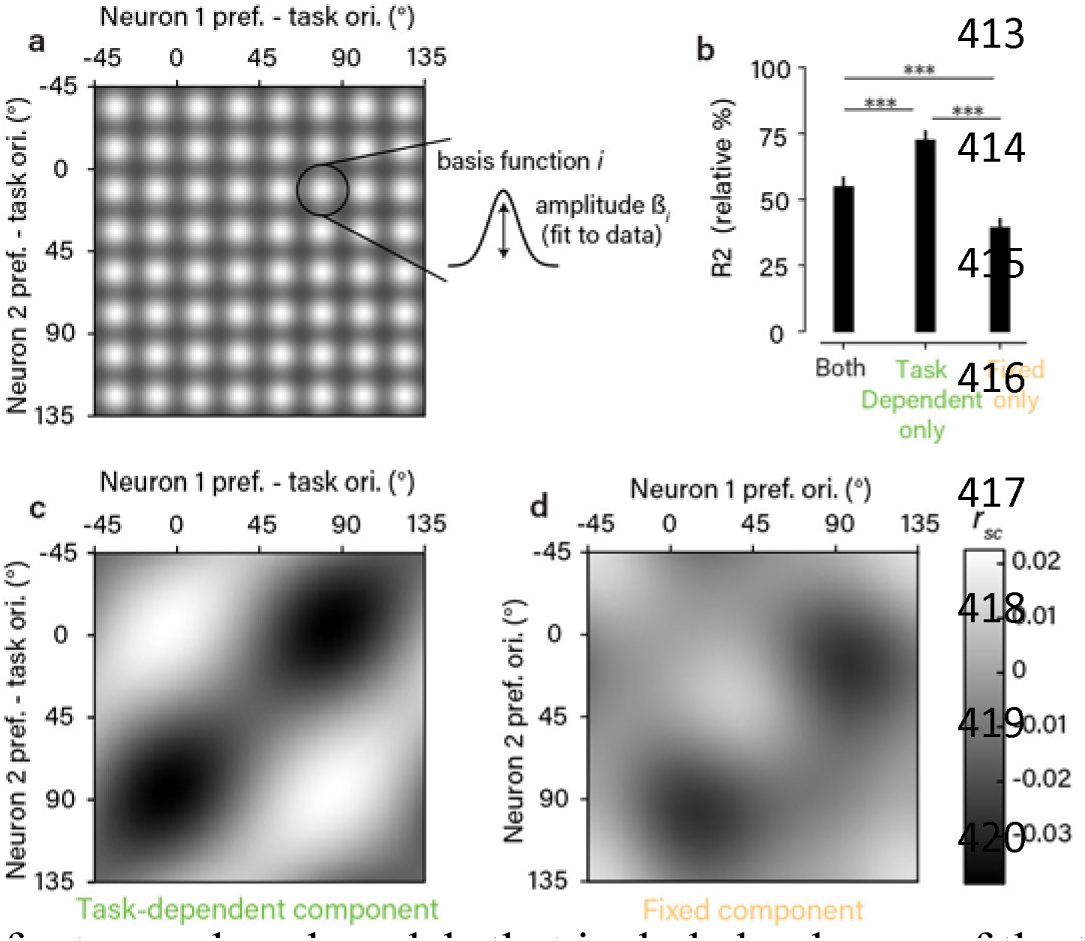
Segregating fixed and task-dependent components of *r*_*sc*_ structure. **a**. Schematic of the regression model used to estimate fixed and task-dependent components of *r*_*sc*_ structure. Each component was a matrix composed of a grid of 8x8 von Mises basis functions, with amplitudes fit to the observed *r*_*sc*_ measurements. **b**. Goodness-of-fit for the model that included both components and for two reduced models that included only one of the two components. Values are expressed relative to an estimate of the explainable variance in the data (see Methods). Error bars are +/- 1 SEM obtained from repeated 50-fold cross-validation. Statistical differences in goodness-of-fit (p<0.001 in all cases) were based on a one-sided test obtained in the same way. **c,d**. Estimated components from the combined model. The amplitude of the task-dependent component (c) was considerably larger than the fixed component (d) by a factor of 2.1 (computed using the varance across the fitted basis function amplitudes), and closely resembled the lattice-like shape of the task-aligned, average *r*_*sc*_ matrix (Fig. 2e). Note that orientation preferences for the task-dependent component are expressed relative to the task orientations. Mean *r*_*sc*_values are close to 0 due to the inclusion of a model constant.

This modeling approach allowed us to address two related questions. First, the form of the fitted components serves to identify the nature of the dynamic and fixed *r*_*sc*_ structure in the V1 population. Second, comparing models that included either or both components provides a quantitative test for the origin of the *r*_*sc*_ structure we observed. When we jointly fit both components to the data, the inferred task-dependent component (Fig. 3c) recapitulated the lattice-like structure we observed in the average data (Fig. 2e). The fixed component (Fig. 3d) was smaller in amplitude and, interestingly, appeared also to contain a weak lattice-like structure, offset by approximately 30º. This is likely due to the fact that we did not uniformly sample across all possible task contexts, with tasks discriminating orientations near 30º/120º being overrepresented (see Fig. 2b). Next, we compared reduced models in which only one of the two components was used. Strikingly, cross-validated model accuracy was increased when we removed the fixed component entirely, but reduced by about half when we removed the task-dependent component (Fig. 3b). This suggests that the dependence of *r*_*sc*_ on orientation preference in our data can be explained as a completely dynamic phenomenon, with no additional dependence that is invariant to the task. We found that that all of these modeling results could be replicated when the fixed and task-dependent components were parametrized in a different way (using a variable number of basis functions with locations fit to the data, instead of a fixed grid of basis functions; see Methods and Supplementary Fig. 8), suggesting the conclusions do not depend on the particular parametric assumptions that were made.

We were interested in the effect of task context on *r*_*sc*_ structure, so it made sense to focus on the dependence of *r_sc_* on orientation preference. However, *r*_*sc*_ depends on a large number of factors irrelevant to the present study, such as physical proximity between pairs and similarity in tuning along many stimulus dimensions apart from orientation^3,22^. This implies that a model that describes the dependency on orientation preference correctly will only explain a small fraction of the variance in *r*_*sc*_. (This can be appreciated in Fig. 2d and f, where pairs with similar locations on the abscicca have substantial variation in *r*_*sc*_.) To estimate this fraction, we assessed the accuracy with which we could predict individual *r*_*sc*_ values from a smoothed matrix built with other pairs. This showed that, in principle, 3.6% of the variance is explainable, of which the majority was explained by the regression model above. We also found that, across cross-validation folds, the fitted model components were highly consistent (mean correlation of 0.99), suggesting the inferred structure is robust to noise in the data despite the low absolute value of variance explained. Additionally, as we will discuss, the task-dependent pattern of *r*_*sc*_ we identify is likely to be critically important during performance of the task despite the low fraction of total variance in *r*_*sc*_ it explains. However, it is important to point out that our data cannot directly speak to the origin of rsc structure in V1 except as it varies as a function of preferred orientation.

### *R_sc_* structure during task performance reflects a single mode of variability

In the modeling discussed so far, we aimed to describe a fixed and task-dependent component of *r*_*sc*_ structure with as few assumptions as possible. Having established that the observed *r*_*sc*_ structure can be best described assuming it is entirely task-dependent, we next sought to identify a more parsimonious and intuitive description of this task-dependency. We started with the observation that the pattern we observed – increased correlation between pairs preferring the same task orientation and decreased correlation for pairs preferring opposing task orientations – would be consistent with feature-selective feedback which varied in its allocation from trial to trial between the two task-relevant orientations, as has been shown in recent theoretical studies^29,34^. To quantify this observation, we performed an eigendecomposition of the smoothed, average *r*_*sc*_ matrix (Fig. 4a). We found that it had a single eigenvalue significantly larger than would be predicted by chance, consistent with the correlation structure being determined largely by a single mode. Moreover, the first eigenvector contained a peak and trough at the two discriminandum orientations, respectively, suggesting a mode of variability which increases the firing rate of neurons supporting one choice and decreases the firing rate of neurons supporting the other choice (Fig. 4b). To model this, we assumed all observed *r*_*sc*_ values could be predicted by a single eigenvector which we constrained to be the difference of two von Mises functions centered 90° apart with variable amplitude and width (see Fig. 4c). We found that this simpler model in fact performed better than the more complex regression model in predicting individual *r*_*sc*_ values, capturing about 80% of the explainable variance in *r*_*sc*_(see Fig. 4e). This suggests that the *r_sc_* structure we observed in V1 could indeed be well described as the result of a single source of covariability that changed dynamically with the task.

**Fig. 4.**
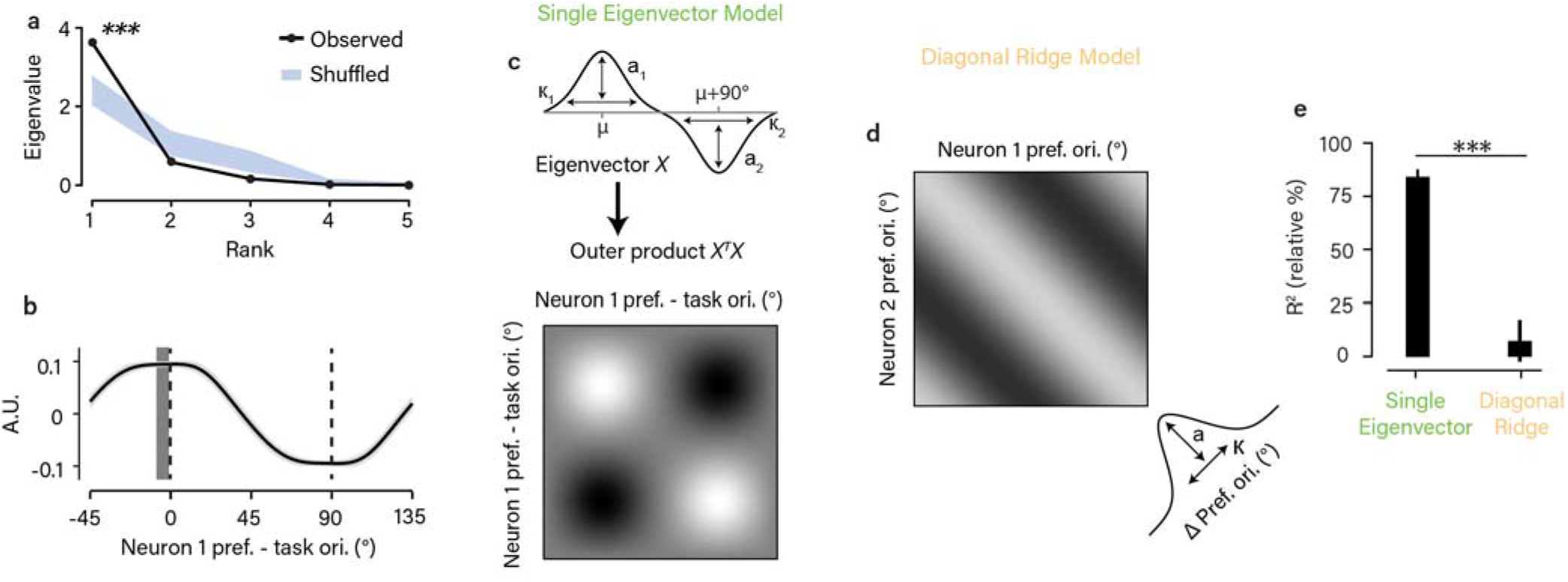
*R_sc_* structure during task performance reflects a single mode of variability. **a**. Eigenspectrum for the average, task-aligned *r*_*sc*_ matrix in Fig. 2e. The largest eigenvalue exceeded chance (p<0.001, permutation test, one-sided). The chance distribution (mean +/- 1 SEM in blue) was determined by adding a random offset to the preferred orientations of each of the 811 pairs (i.e. permuting each *r*_*sc*_ value along the diagonal). **b.** The eigenvector corresponding to the largest eigenvalue in (a). We first removed the mean *r*_*sc*_ value from the matrices to ignore any flat eigenvectors. Error bar is +/- bootstrap SEM. The dark gray vertical bar indicates the peak in the eigenvector +/- 1 bootstrap SEM. This was not significantly different from 0º (p=0.078, bootstrap test, one-sided), indicating close alignment with the task. **c.**Schematic of “single eigenvector” model, in which *r*_*sc*_ structure is described as the outer product of a vector parameterized as the difference between two von Mises functions 90° apart. **d**. Schematic of the “diagonal ridge” model in which *r*_*sc*_ structure depended only on the difference in preferred orientations, independent of task. This dependence was modeled as a von Mises function centered on zero. **e**. Goodness-of-fit for the models in (c) and (d), calculated as normalized % variance explained, as in Fig. 3. Error bars around the mean and statistical comparison between models obtained through repeated 50-fold cross-validation of the set of 811 pairs.

We compared the “single eigenvector” model with another simple model that more closely reflected standard assumptions about *r*_*sc*_ structure in sensory brain areas. This model predicted that *r*_*sc*_depends only on the difference in preferred orientation between pairs of neurons regardless of task^5,9,10^(“limited-range correlations” yielding an *r*_*sc*_ matrix with a diagonal ridge) and would be consistent with *r*_*sc*_ structure due to common afferent inputs. We modeled this dependence as a von Mises function of preferred orientation difference (Fig. 4d). This model performed much worse in predicting the observed set of *r*_*sc*_ values, in fact not exceeding chance performance (Fig. 4e). (This qualitative difference in model performance was replicated in both subjects individually; see Supplementary Fig. 2). Importantly, both of these simple models predict a dependence of *r*_*sc*_ on preferred orientation difference similar to what we found in the data (Fig. 2d) and has been observed previously^8,20–23^ – however, in the case of the “single eigenvector” model, this is due to task-dependent changes in *r*_*sc*_ while for the “diagonal ridge” model, there is no effect of task context. Notably, we found that during the passive fixation blocks, the “diagonal ridge” model performed better (Supplementary Fig. 3c), quantitively supporting the observation that the task-dependent correlations we observed require active task engagement.

### Effect of task-dependent *r*_*sc*_ structure on neural coding

We next sought to address the functional importance of the *r*_*sc*_ we observed on sensory coding. Many studies have shown that *r_sc_* in sensory neurons can decrease the sensory information that can be decoded, particularly when *r*_*sc*_ resembles correlations due to task-related stimulus changes^5,8–15^. We estimated this task-related stimulus correlation as the product of the slopes of a pair’s mean response functions along the task axis (i.e. as a function of orientation signal strength; Fig. 5a)^16^, normalized by the product of the neuronal variances. When we plotted these values as a smooth, task-aligned matrix (Fig. 5b), we observed a lattice-like pattern strikingly similar to the observed *r*_*sc*_ matrix (Fig. 2e). Confirming this similarity, the task-dependent component of *r_sc_* structure identified by the regression model was highly correlated on a pair-by-pair basis with the stimulus-induced correlations (*r*=0.61, Fig. 5c). This matches our earlier observation that *r*_*sc*_ structure was consistent with feedback that alternatingly targeted the task-relevant neuronal pools, which is similar to the effect of varying the stimulus along the axis defining the task.

**Fig. 5.**
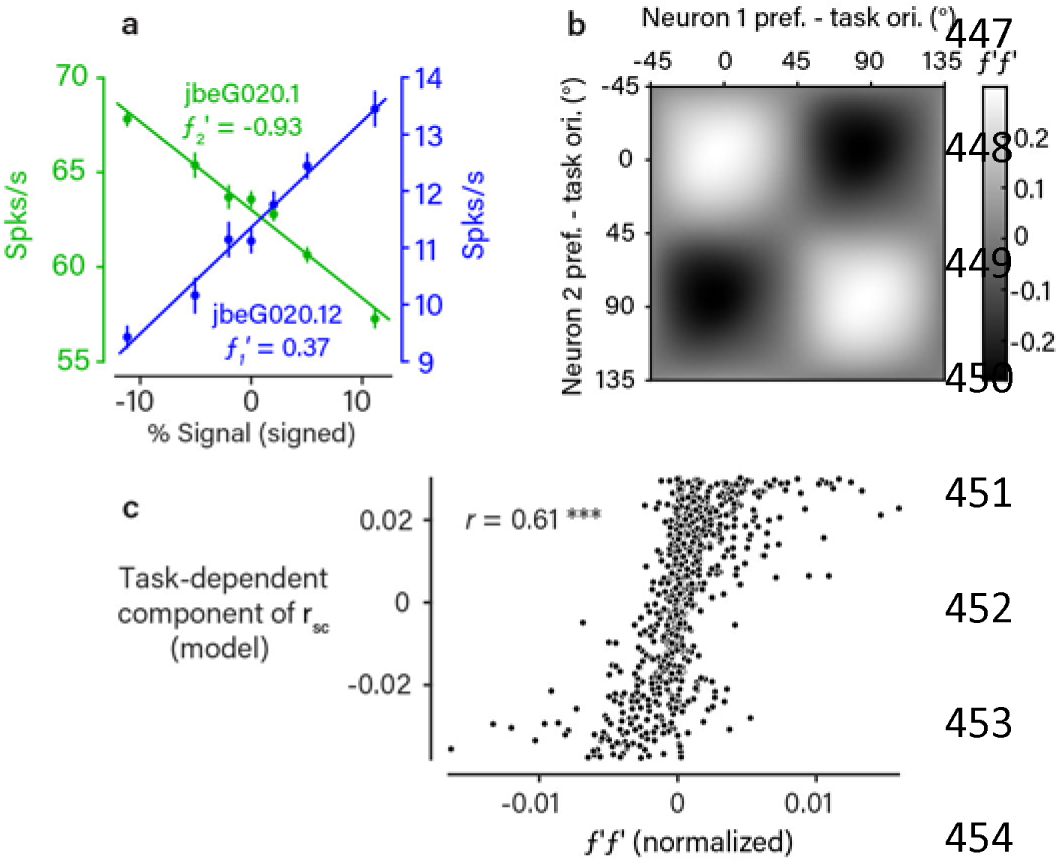
*R_sc_* structure matches effect of task-related stimulus variability. **a**. Responses (mean +/- 1 SEM, n=1,049 trials) to the stimuli used in the task at various signal strengths for two example neurons. For the purposes of illustration, the two task orientations are simply labeled positive and negative. This pair was typical in that the response functions (*f*_*1*_ and *f*_*2*_) are approximately linear over the range of signal strengths used. For this reason, we calculated the response correlation introduced by tuning similarity as the normalized product of the derivatives *f*_*1*_◻*f*_*2*_◻ ^16^. **b**. The matrix of *f***D***f***C** values, as a function of task-aligned pairwise orientation preference, obtained using kernel smoothing as in Fig. 2. We observed a pattern that was very similar to the structure of *r*_*sc*_ we observed during task performance (Fig. 2e). **c**. Scatter plot of the task-dependent (putatively top-down) component of *r_sc_* (Fig. 3 c) against normalized *f*◻*f*◻ values for each recorded neuronal pair. The two were highly correlated across the population (Pearson’s r=0.62, p<0.001, bootstrap test, one-sided).

Thus, the observed *r*_*sc*_ structure appears not to improve, but rather to degrade, the sensory representation. However, our results highlight a problem with this interpretation and any purely feedforward account of the functional role of *r*_*sc*_. Namely, *r*_*sc*_ that is generated endogenously need not be problematic at all (e.g. if the decoder had access to those endogenous signals). Indeed, the propagation of feedback signals that are matched to the statistics of the relevant sensory stimuli may be an adaptive strategy for bringing prior knowledge to bear, as predicted by recent models of probabilistic perceptual inference^28,29^. *R*_*sc*_ resembling stimulus-induced correlations emerge in such models^28^ as a consequence of the subject developing the appropriate priors about the task, yielding predictions that both match our empirical findings and offer a normative explanation.

### Relationship between *r*_*sc*_ structure and perceptual choice

Correlations between trial-to-trial variability of single neurons and choice^35,36^ have been frequently observed throughout sensory cortex. Theoretical studies have emphasized that this suggests the presence of spike-count correlation with a particular structure^17–19,36,37^. After all, if many sensory neurons have variability that is correlated with choice, this implies that neurons supporting the same choice are themselves correlated. However, this could be compatible with either or both of two causal mechanisms: 1) correlated fluctuations directly affect the choices a subject makes trial to trial (a feedforward source of choice-related activity); or 2) the correlated fluctuations reflect variation across trials in a feedback signal related to the upcoming choice (a feedback source). As we show, our detailed measures of *r*_*sc*_ structure during task performance can address this ongoing debate.

First, we reasoned that a signature of feedback related to the upcoming choice would be *r*_*sc*_ structure in V1 whose magnitude depends systematically on variability in choice. Consistent with this prediction, we found that the amplitude of the *r*_*sc*_ structure was attenuated on high-signal trials relative to 0% signal trials, in a manner which depended systematically on signal strength (Fig. 6a,b). However, this attenuation was modest, even at the highest signal level we analyzed (11% reduction), despite the highly uneven distribution of choices. This rules out the extreme scenario in which feedback perfectly reflects choice. Supporting this conclusion, we found that the *r*_*sc*_ structure, when calculated using only spikes from different 200-ms windows during the trial, showed a stable timecourse (after a precipitous drop at the first time point) and did not grow in amplitude with decision formation (Fig. 7). Taken together, these observations support the conclusion that the *r*_*sc*_ structure reflects variation in feedback signals only partially correlated with the subject’s final choices. These could include a combination of bias, attention to orientation, prior beliefs, and/or a decision variable.

**Fig. 6.**
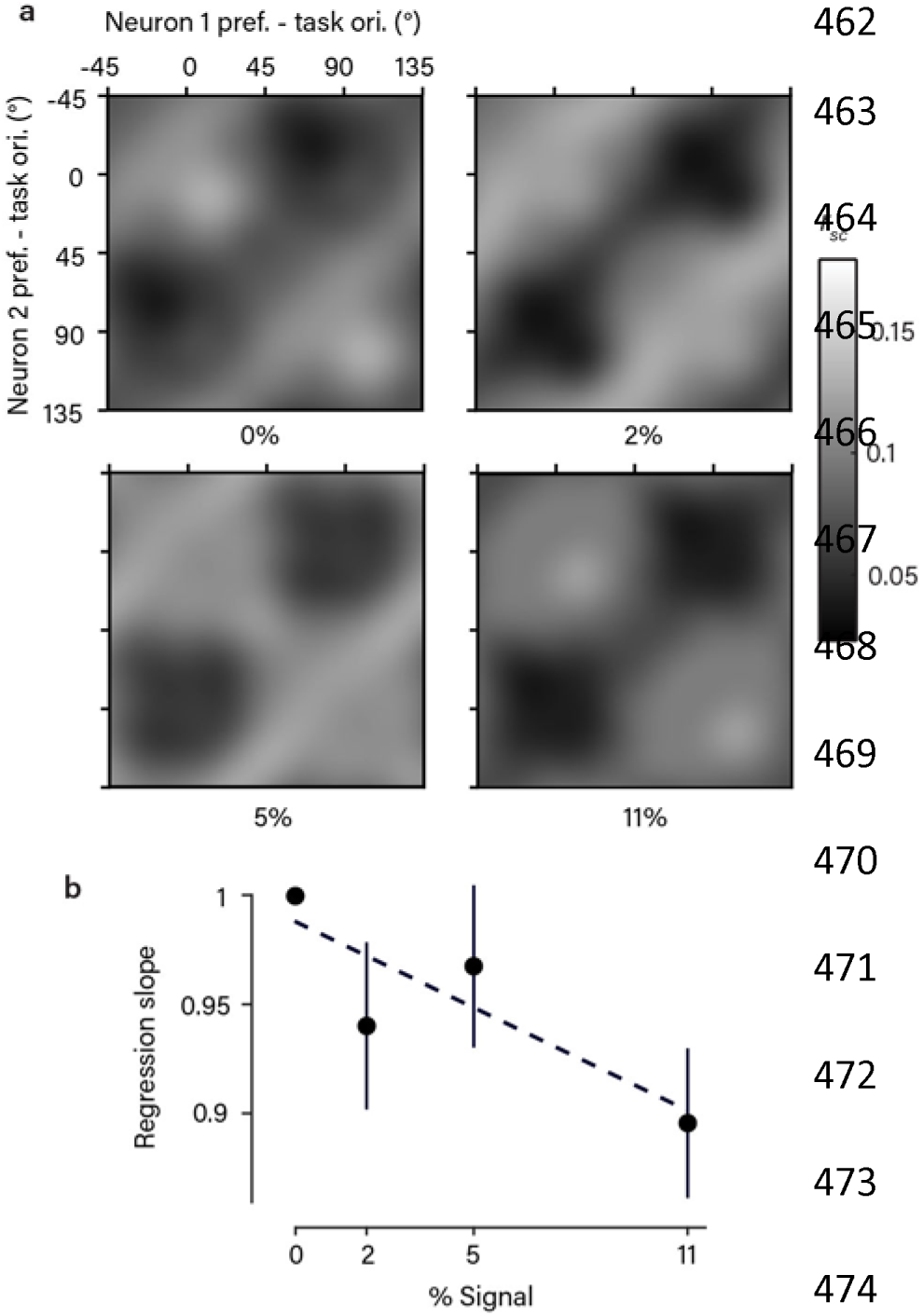
*R_sc_* structure depends on variability in choice. **a**. The average, task-aligned *r*_*sc*_ matrix (as in Fig. 2e), shown separately for each stimulus strength. Note that 0% signal trials involved statistically identical stimuli across all task contexts. A qualitatively similar structure was apparent at nonzero signal levels. (Spike counts were z-scored to eliminate the effect of stimulus drive; see Methods). **b.** Scatter plot showing the slope of a regression line comparing the *r*_*sc*_ values measured at each signal level against the *r*_*sc*_ values measured at the 0% signal level. This quantity indicates the degree of attenuation of the *r*_*sc*_ structure at a given signal level. We observed a weak but significant negative correlation (Pearsons’s *r*, p=0.038, bootstrap test, one-sided) with signal strength (error bars are +/- 1 bootstrap SEM around the mean of the 811 pairs), implying the *r*_*sc*_ structure is attenuated on high-signal trials, when there was also less variability in choice. Dotted line is fitted regression line.

**Fig. 7.**
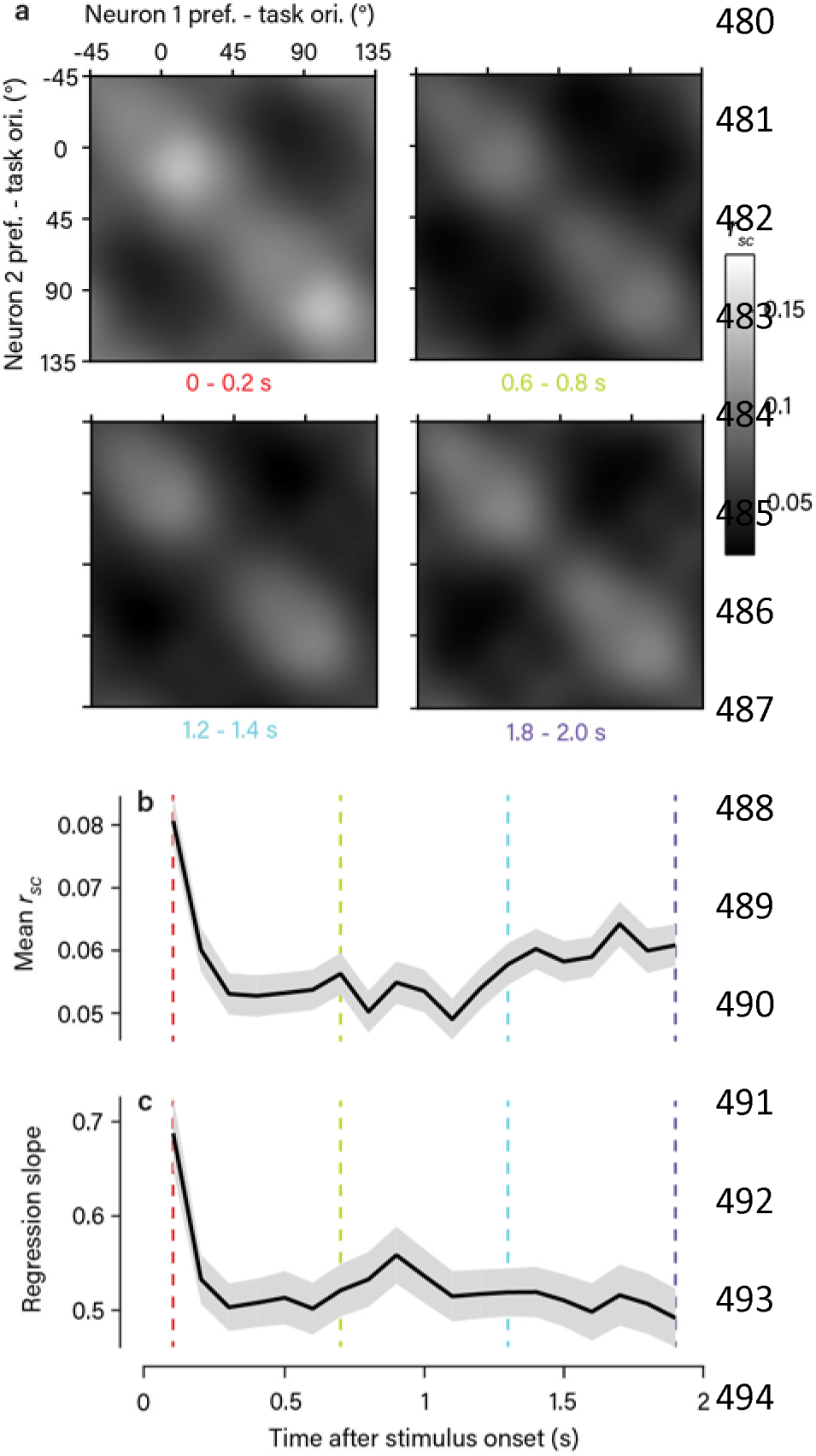
Temporal dynamics of *r*_*sc*_ structure. **a.** The average, task-aligned *r*_*sc*_matrix (as in Fig. 2e) obtained using spike counts from 200-ms windows during the stimulus presentation. A similar structure was present at all time points (4 examples shown). **b-c**. Plots showing the temporal dynamics of two statistical measures of the observed *r*_*sc*_ structure (mean +/- 1 bootstrap SEM). The colored lines indicate the example time points shown in (a). The population mean *r*_*sc*_ value (b) showed a sharp drop shortly after stimulus onset, as seen in other studies^50^, and then a gradual recovery over the course of the trial. The amplitude of the *r*_*sc*_ structure, quantified using the slope of the regression line of *r*_*sc*_ obtained in each 200-ms window against *r*_*sc*_ obtained from trial-length spike counts, is in (c). Apart from an increase at the first time point, likely due to the onset of the visual stimulus, this showed no significant modulation over the course of the trial. Note that values are all significantly less than 1 because smaller counting windows introduced a source of uncorrelated noise across trials.

Next, we assumed standard feedforward pooling (i.e. linear readout weights applied to the sensory pool) to determine if the observed *r*_*sc*_ structure would be quantitatively consistent with the observed choice-related activity. To do this, we made use of recent theoretical work which analytically relates *r*_*sc*_ structure, readout weights, and choice-related activity^17^. We calculated Choice Probability (CP), which quantifies the probability with which an ideal observer could correctly predict the subject’s choices using just that neuron’s responses^35,36^, for each recorded neuron. We found an average CP of 0.54 for task-relevant neurons, significantly above chance level (Fig. 8a) and similar in magnitude to another study using the same task^31^. We found that the *r*_*sc*_ structure we observed would be sufficient to produce a pattern of CP across the population consistent with the data (Fig. 8b,c), across a wide range of possible readout schemes (Supplementary Fig. 9). Next, we considered the contribution of the different inferred sources of *r*_*sc*_ to CP. (For top-down sources of correlation this is equivalent to assuming that the sensory population is read out without taking into account the top-down signal.) This allows us to treat all sources of *r*_*sc*_ equivalently, and compare them quantitatively. When we considered a population containing only the “task-dependent” component of *r*_*sc*_ structure identified in the regression model (Fig. 3c), predicted CP was only slightly reduced. Assuming only the “fixed” component (Fig. 3d), however, drastically reduced predicted CP below what we observed (Fig. 8b,c). Thus, our data rule out the view that a significant component of CP merely reflects the feedforward effect of stochastic noise in the afferent sensory pathway. Instead, the main feedforward source of CP appears to depend on task-dependent changes in *r*_*sc*_ structure that subsequently influence perceptual judgments.

**Fig. 8.**
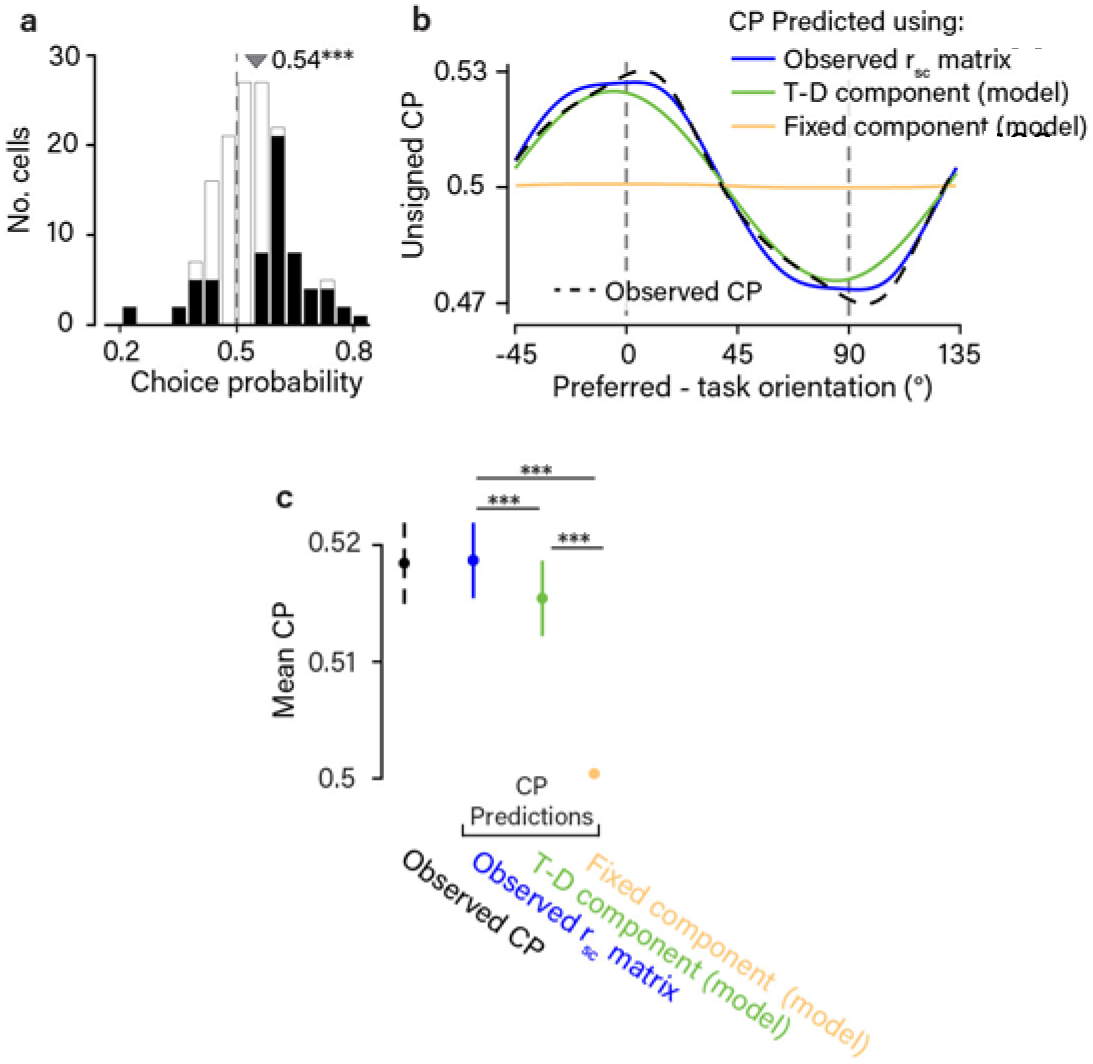
The task-dependent component of *r*_*sc*_ structure accounts for choice-related activity. **a**. Histogram of observed CPs, from the subset of neurons (*n*=144) significantly preferring one of the two task orientations (*d*◻>0.9 at highest signal level). Mean CP of 0.54 exceeded chance (p<0.001, bootstrap test using cell resampling, one-sided). CPs that were individually significant (p<0.05, bootstrap test using trial resmpling, one-sided) are shown in black. **b**. We tested the known analytical relationship between spike-count correlations, readout weights, and CPs, under the assumption of a linear decoder applied to a population of sensory neurons^17^ (see Methods). Here CP is defined as a continuous function of task-aligned preferred orientation, analogous to our description of the *r_sc_* matrix in Fig. 2e. The dashed black line shows the profile of CP observed across preferred orientations, after smoothing with a von Mises kernel approximating a wrapped Gaussian with 10° s.d. We applied a fixed sign convention to the CP values across all neurons, equivalent to arbitrarily calling the 0°-choice the preferred one. The predicted CP profiles (solid lines) show the CP elicited by reading out a sensory population with different *r*_*sc*_ structures. Readout weights across orientations were unobserved and the profiles shown are averages of a large set generated from different assumed readout weight profiles (see Methods). **c**. Mean CP (using the traditional sign convention) associated with the profiles in (b), +/- 1 bootstrap SEM obtained by cell resampling (n=811 neurons). Note that the mean CP shown here is different to the one shown in (a) because all neurons are included, regardless of their orientation preference

## Discussion

Spike-count correlations between sensory neurons have typically been described as reflecting noise that corrupts sensory encoding^5,8–15^. However, little is known about the origin of *r*_*sc*_, and it may instead be due to changes over time in feedback signals. We addressed this by recording from populations of V1 neurons using multi-electrode arrays while macaque subjects performed a set of orientation discrimination tasks. This approach allowed us to estimate the entire matrix describing the dependence of *r*_*sc*_ on pairwise orientation preference (Fig. 2), providing an unprecedently clear picture of *r*_*sc*_ structure in a behaving animal. By determining to what extent the *r*_*sc*_ matrix was fixed, and what extent it changed with task, we could infer the relative importance of feedforward and feedback pathways in generating it (Fig. 3). We found systematic and novel structure in the *r*_*sc*_ matrix that changed in a predictable manner with the task. Using multiple modeling approaches, we found that the fixed *r*_*sc*_ structure was much smaller than the task-dependent structure, so much so that we could not estimate a fixed component reliably. Remarkably, a single source of task-dependent feedback captured the pattern we observed (Fig. 4). This feedback input increased and decreased the firing rate of neurons tuned for the two task-relevant orientations in a push-pull manner.

Our results suggest the possibility that variability in feedback is a major source of *r*_*sc*_ structure in sensory cortex. The role of feedback may be even more pronounced in areas downstream of V1 which typically show a greater degree of extra-sensory modulation^31,38–40^. At the same time, we cannot rule out a larger role of feedforward inputs in generating patterns of *r*_*sc*_ defined in different ways than those uncovered here. For example, because our measures of *r*_*sc*_ structure involved smoothing, we cannot rule out the possibility that the fine-grained structure of *r*_*sc*_ behaves in ways not captured by our analysis.

Our results are consistent with, and expand upon, a prior study that also measured task-dependent changes in *r*_*sc*_^27^. In that study, single pairs of direction-selective MT neurons were recorded while subjects performed two direction discrimination tasks chosen by the experiments to probe the effect of task context: one in which the neurons contributed to the same choice (“same-pool condition”) and one in which they contributed to opposite choices (“opposite-pool condition). This amounts to a selective sub-sampling of the *r*_*sc*_ structure. While this identified some degree of task-dependence, the implications remained unclear. By contrast, the present study involved recordings from large simultaneously recorded populations, which achieved much better coverage of the full *r*_*sc*_ structure. This revealed the detailed structure of the task-dependence and provided the basis for quantitative modeling and novel conclusions. For the purposes of comparison, we plotted our data in an analogous way to the prior study and found qualitatively similar results (Supplementary Fig. 10).

Consistent with several past studies^30,41,42^, we found evidence for choice-related feedback, as shown by the finding that correlated fluctuations in V1 are more pronounced on trials where the subject’s choices were more variable (Fig. 6). However, this effect was relatively weak, and we observed that task-dependent *r*_*sc*_ structure did not grow in amplitude with decision formation (Fig. 7), suggesting processes indirectly related to choice may be responsible for the feedback generating the correlations. More importantly, we found that the standard assumption that correlated fluctuations influence choice through feedforward pathways^17–19,36,37^ predicted CP in the V1 population that matched the data (Fig. 8), the first empirical test of the theoretical relationship between *r*_*sc*_ in sensory neurons, CP, and readout^17^. However, the *r*_*sc*_ structure responsible changed with the task, demonstrating that it does not simply reflect afferent noise. Taken together, our results instead favor the notion that choice-related activity comes about through self-reinforcing loops of reciprocal connectivity between cortical areas, as has also been suggested by other studies^29,42,43^.

The task-dependent modulation of *r*_*sc*_ we observed did not appear to be beneficial to task performance (Fig. 5), at least not in the manner this has typically been examined (i.e. feedforward decoding of the sensory population alone). Instead, the inferred feedback signals appeared to mimic task-relevant stimulus changes, confounding the choices of an observer using only the sensory population. However, because the correlations reflect downstream computations, they need to not be limiting in this way to the subject. Thus our results highlight the fundamental insufficiency of considering the theoretical implications of *r*_*sc*_ in terms of purely feedforward frameworks, as almost all such studies have done to date.

The inferred source of task-dependent feedback resembles previous reports about the effects of feature-based attention on visual cortical neurons^34,44^. Feature-based attention enhances the firing rate of neurons tuned for the attended stimulus feature, and decreases the firing rate of neurons tuned for unattended stimulus features. One possibility is that our task engages feature-based attention which varies over time in its allocation between the two task-relevant orientations. This does not appear to provide an adaptive increase in the amount of relevant stimulus information encoded, contrary to traditional descriptions of attention^45,46^. However, as discussed above, once a top-down contribution to correlations is recognized, it is not possible to infer the amount of sensory information available to a decoder from the activity of a population of sensory neurons alone.

Our findings thus emphasize the need for new normative models that predict context-dependent feedback during perceptual processing. Currently, models based on hierarchical probabilistic inference^28,29,47^ do predict such feedback signals, and account for many of our experimental findings. This work builds on the longstanding idea that the goal of a perceptual system is to generate valid inferences about the structure of the outside world, rather than to faithfully represent sensory input^48,49^. This requires combining sensory input with prior beliefs, both of which can introduce correlated variability. During perceptual decision making, correlations resembling those induced by the stimulus naturally emerge as a consequence of the subject acquiring the appropriate prior beliefs about the structure of the sensory environment^28^. Clearly, further development of this and other models of perceptual processing are needed to generate quantitative predictions which can be further tested empirically.

## Acknowledgements

We thank Bob Wurtz and James McFarland for useful discussions; Richard Krauzlis, Bevil Conway, and Ali Ghazizadeh for comments on an earlier version of the manuscript; and Beth Nagy, Irina Bunea, and Denise Parker for veterinary care.

## Author Contribution

A.G.B. and B.G.C. conceived and designed the experiments. A.G.B. performed the experiments and all aspects of the analysis. A.G.B. and B.G.C. wrote the paper. R.H. advised and assisted with the data analysis and the paper. B.G.C. advised at all stages.

## Methods

### Electrophysiology

We recorded extracellular spiking activity from populations of V1 neurons in two male, awake, head-fixed rhesus monkeys (*Macaca mulatta*). For the majority of the recordings, monkey ‘*lem*’ was 14 while monkey ‘jbe’ was 16 years old, before which time they had each experienced extensive behavioral training, including on other behavioral paradigms for monkey ‘*lem*’. Monkey ‘*lem*’could not be pair housed due to antisocial behavior. Both monkeys were implanted with a head post and scleral search coils under general anaesthesia^51^. In monkey ‘*lem*’, a recording chamber was implanted over a craniotomy above the right occipital operculum, as described previously^52^, by which we introduced linear microelectrode arrays (U and V-probes, Plexon; 24-contacts, 50 or 60 µm spacing) at an angle approximately perpendicular to the cortical surface with a custom micro-drive. We positioned the linear arrays so that we roughly spanned the cortical sheet, as confirmed with current-source density analysis, and removed them after each recording session. In monkey ‘*jbe*’, a planar “Utah” array (Blackrock Microsystems; 96 electrodes 1mm in length inserted to target supragranular layers, 400 um spacing) was chronically implanted, also over the right occipital operculum. All procedures were performed in accordance with the U.S. Public Health Service Policy on the humane care and use of laboratory animals and all protocols were approved by the National Eye Institute Animal Care and Use Committee.

Broadband signals were digitized at 30 or 40 kHz and stored to disk. Spike sorting was performed offline using custom software in MATLAB^®^. First, spikes were detected using a voltage threshold applied to high-pass filtered signals. Next, triggered waveforms were projected into spaces defined either by principal components or similarity to a template. Clusters boundaries were finally estimated with a Gaussian mixture model, and then rigorously verified and adjusted by hand when needed. In the linear array recordings, spike sorting yield and quality was substantially improved by treating sets of three or four neighboring contacts as “n-trodes”. As this was not possible with the Utah array due to the greater interelectrode spacing, we excluded pairs of neurons recorded on the same electrode to avoid ontamination by misclassification. Neurons from separate recording sessions were treated as independent. To reduce the possibility that a single neuron from the Utah array contributed to two datasets, we included only sessions that were separated by at least 48 hours (with a median separation of 5 days). We excluded from analysis those neurons whose mean evoked firing rate did not exceed 7 spikes/second.

### Visual stimuli

All stimuli were presented binocularly on two gamma-corrected cathode ray tube (CRT) monitors viewed through a mirror haploscope, at 85 or 100Hz. The monitors subtended 24.1° x 19.3° of visual angle (1280 x 1024 pixels). The stimuli presented during performance of the discrimination task 31 consisted of bandpass filtered dynamic white noise, as described previously. Briefly, stimuli were filtered in the Fourier domain with a polar-separable Gaussian. The peak spatial frequency was optimized for the recorded neuronal population (1 and 4 cpd medians for ‘*lem*’and ‘*jbe*’,respectively) while the peak orientation could take one of two orthogonal values the animal had to discriminate in a given session. The angular s.d. of the filter modulated the orientation bandwidth and was varied trial to trial. A 2D Gaussian contrast envelope was applied to the stimulus so that its spatial extent was as small as possible while still covering the minimum response fields of the neuronal populations being recorded. The median envelope s.d. was 0.6 degrees for both animals. The median stimulus eccentricity was 5.4 degrees for ‘*lem*’and 0.5 degrees for ‘*jbe*’. In Fig. 1, we quantify orientation bandwidth as % signal strength. This was calculated as 100 * R,where Ris the length of the resultant vector associated with the angular component of the stimulus filter. To perform psychophysical reverse correlation (PRC) for orientation (Supplementary Fig. 1), we summarized the orientation energy of the stimulus on each trial as the radial sum of its 2D amplitude spectrum (averaged across frames) to remove information about spatial frequency and phase.

We estimated neuronal orientation preferences in separate blocks of trials, using 420-ms presentations of the following types of stimuli, presented at a range of orientations: 1) an orientation narrowband version of the stimulus described above (10° angular s.d.); 2) sinusoidal gratings; and 3) circular patches of dynamic 1D noise patterns (random lines). The preferred orientation of a neuron was calculated as the circular mean of its orientation tuning curve. For each neuron, from among the set of tuning curves elicited by the different stimulus types described above, we chose as the final estimate of preferred orientation the one with the smallest standard error, obtained by resampling trials. We excluded from further analysis all neurons where this exceeded 5°. On a subset of sessions, we also used these orientation-tuning blocks to present examples of the 0%-signal orientation-filtered noise stimuli. These were presented at the same location and size as during task performance, allowing us to calculate *r*_*sc*_ structure in the absence of task engagement but with identical retinal input.

### Orthogonal orientation discrimination task

The animals performed a coarse orientation discrimination task using the orientation-filtered noise stimuli, as described previously^31^. To initiate a trial, the subject had to acquire a central fixation square. After a delay of 50 ms, the stimulus appeared for a fixed duration of 2 seconds. The trial was aborted if the subject broke fixation at any point during the stimulus presentation, defined as either 1) making a microsaccade covering a distance greater than a predefined threshold (typically 0.5°) or 2) a deviation in mean eye position from the center of the fixation point of more than a predefined threshold, typically 0.7°. At the end of the stimulus presentation, two choice targets appeared. These were Gabor patches of 2-3° in spatial extent, oriented at each of the two discriminandum orientations. The locations of the choice targets depended on the task. For orientation pairs near horizontal and vertical (-22.5° – +22.5° and 67.5° – 112.5°), the choice targets were positioned along the vertical meridian, at an eccentricity of about 3°, with the more vertically-oriented target appearing always in the upper hemifield. For orientation pairs near the obliques (22.5° – 67.5° and 112.5° – 157.5°), the choice targets were positioned along the horizontal meridian, at the same range of eccentricities, with the smaller of the two orientations always appearing in the left hemifield. (We use the convention that horizontal is 0° and that orientation increases with clockwise rotation.) To penalize random guessing, the volume of liquid reward delivered after correct choices was doubled with each consecutive correct choice, up to a maximum of four times the initial amount. Since we were primarily interested in the effect of task engagement on neuronal activity, we applied a behavioral criterion to our data, excluding sessions where the subject’s psychophysical threshold (defined as the signal level eliciting 75% correct performance) exceeded 14% signal.

To determine the influence on *r*_*sc*_ of random fluctuations in the stimulus introduced by the use of white noise, we used a double-pass experimental design^53^ in which each exact stimulus sequence was presented on two separate trials. We calculated the stimulus-induced *r*_*sc*_ for each pair, as described below, after permuting the indices of the paired repeat trials for one neuron’s trial sequence. This eliminated the temporal alignment of the two trial sequences, abolishing stimulus-independent covariability, while preserving the identity between the stimuli associated with the two trial sequences.

We attempted to use as wide a range of task contexts as possible over the course of data collection from both animals, but task contexts were not presented in a randomized way to the subjects, since performing a new task context required several days of retraining. Additionally, data collection and analysis was not performed blind to the experimental conditions – in particular, experimenters were aware what the instructed task context was. For further detailed information on experimental design and reagents, see the Life Sciences Reporting Summary included online.

### Spike-count correlation measurements

Spike-count correlations were calculated as the Pearson correlation between spike counts, counted over the entire duration of the stimulus, with a 50-ms delay to account for the typical V1 response latency. Spike counts were first z-scored separately within each experimental block (typically a set of 100-200 trials lasting about 10 minutes) and each stimulus condition. This removed correlations related to long-term firing rate nonstationarities and allowed us to combine trials at different signal levels without introducing correlations related to similarity in stimulus preference. We used a balanced z-scoring method proposed recently to prevent bias related to differences in choice distributions across signal levels. We excluded pairs that were not simultaneously isolated for at least 25 trials total. The median number of trials per pair during task performance was 752.

Despite the use of z-scoring, any influence of stimulus history on firing rates could in principle introduce a source of covariability that depended on the task context, since the set of stimuli used was not identical across task contexts (only the 0%-signal condition was identical). We ruled out this confound by adapting the z-scoring procedure described above to further remove any information about the preceding stimulus contained in the spike rate on the current trial. To do this, we z-scored spike counts separately within groups of trials for which the current stimulus *and* the stimulus on the preceding trial were the same. This produced identical results to those shown in the main analysis (Supplementary Fig. 5).

A main goal of the study was to measure how spike-count correlation varies with pairwise orientation. We illustrate this dependence in several figures as a smoothed function estimated from measures of *r*_*sc*_ combined across multiple recording sessions, which we then sampled discretely with 1° resolution. The smoothed estimates were obtained using a bivariate von Mises smoothing kernel. A point in the correlation matrix **C** was given as:

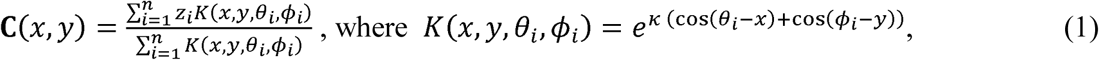

*z*_*i*_ is the *i*^*th*^ *r*_*sc*_ measurement, *θ*_*i*_ and *ϕ*_*i*_ are the preferred orientations of the *i*^*th*^ pair, and Kis the von Mises width parameter. We set K —1.3π, yielding a smoothing kernel closely approximating a bivariate wrapped Gaussian with 15° s.d. (Note that this smoothing procedure was only performed to generate figures in the manuscript, and was not applied as a pre-processing step in any of the quantitative analyses.) In some cases, we expressed the *r*_*sc*_ matrix in a task-aligned coordinate frame (e.g. Fig. 2e), for which the preferred orientations of the *i*^*th*^ pair relative to the task orientations were used for *θ*_*i*_ and *ϕ*_*i*_. Since there were always two orthogonal task orientations, we averaged across both possible alignments, such that C(*x, y*)—C(*x*+ 90°,*y*+ 90°).All angular quantities were doubled for the calculations, as orientation has a period of 180°. To generate the kernel-smoothed profile of CP (Fig. 8), we used a one-dimensional equivalent of the procedure above, in which preferred orientations were parameterized only by a single parameter.

We considered using covariance instead of correlation to measure the covariability of neuronal pairs. However, a key advantage of correlation is that it is insensitive to the variance of the spike counts. By contrast, measures that do not normalize for spike-count variance will effectively overweight more variable pairs in any population analysis. In addition, using spike-count correlation allowed us to combine z-scored counts across stimulus conditions. This substantially increased the signal-to-noise ratio of our measurements. As a confirmation that this approach yielded results that generalize, we measured the average, task-aligned spike-count covariance matrix, using the same approach as we used to generate the *r*_*sc*_ matrix in Fig. 2e. To estimate the spike-count covariance between a given pair of neurons without including an effect of common stimulus drive, we used an average of the covariance values measured separately for each stimulus condition, weighted by number of trials. We found that the pattern in the spike-count covariance matrix was closely similar to the *r*_*sc*_ matrix (Supplementary Fig. 11). This confirms that our main results are not dependent on the use of *r*_*sc*_ measured with normalized spike counts.

### Regression model

We used a multilinear regression model to identify fixed and task-dependent components of the structured correlations we observed. We describe the set of observations (811 individual pairwise *r*_*sc*_ measurements) in terms of a set of two underlying correlation structures: one defining *r*_*sc*_ as a function of pairwise preferred orientation alone (“fixed”) and the other defining *r*_*sc*_ as a function of pairwise preferred orientation relative to the task orientations (“task-dependent”). In order to provide a continuous and smooth description of the data, each component was parameterized as the sum of an array of *n x n* evenly spaced basis functions. Each observation, *y*_*i*_, was expressed as:

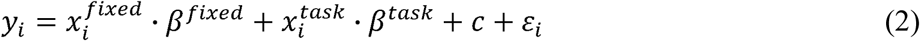

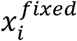and 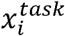are length-n^2^ vectors of loadings onto the basis functions, which were given by evaluating the basis functions at the location corresponding to the pairwise orientation preference of the *i*^*th*^ pair. *β*^*fixed*^ and *β*^*task*^ are the length-n vectors of amplitudes of the basis functions (coefficients to be fit), cis a model constant, and · is the element-wise product. For the basis functions, we used bivariate von Mises functions, with no correlation and equal width in both dimensions. Thus the *k*^*th*^ loading 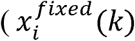or 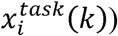was given by:

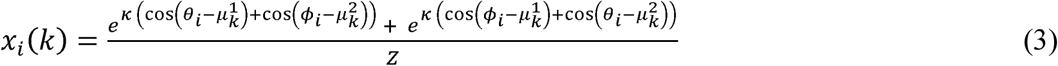

where *θ*_*i*_ and *ϕ*_*i*_ are the preferred orientations of the *i*^*th*^ pair (relative to the task orientations in the case of the task-dependent loadings), *μ*_*k*_ is a pair of orientations defining the location of the *k*^*th*^ basis function, Zis a normalization constant such that the sum of all loadings for observation *i* 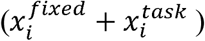 is 1, and Kis the basis function width. Two terms are needed to express the loadings because the data are correlations: the first term describes the upper triangular portion and the second describes the lower triangular portion. Again, angular quantities were doubled. Kacts as a smoothing hyperparameter. We found that arrays of 8x8 were sufficient to describe the structure of the two components. It was sufficient only to fit the upper triangular portion of the array of basis functions. Thus, each component was described by 36 parameters (although the effective number of parameters is significantly less because of the basis function smoothness and the ridge penalty). We fit the model using ridge regression. The unique optimal solution could therefore be derived analytically as 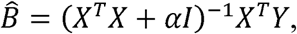where X is the concatenated design matrix combining *x*^*flxed*^ and *x*^*task*^ and *α* is the ridge parameter, which penalized the squared amplitude of the basis functions. The optimal values of the hyperparameters *α* and *κ* were chosen under 50-fold cross-validation.

To ensure our results were not due to the particular way the above model was constructed, we compared them to those obtained using a conceptually similar regression model. In this alternative model, instead of a grid of basis functions with fixed locations, we allowed each component to be described as the sum of a variable number of von Mises basis functions with locations (as well as width and amplitude) fit to the data, again using least squares. This alternative model allowed, in principle, for fewer parameters and for fine details in the *r*_*sc*_ structure to be captured by allowing some basis functions to have small width. The relative contribution of the fixed and task-dependent components of *r*_*sc*_ structure could be tested in terms of the number of basis functions needed to best explain the data. In this case, the *k*^*th*^ loading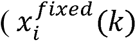or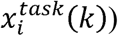 was given by:

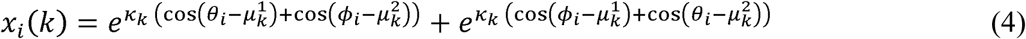

where *θ*_*i*_ and *ϕ*_*i*_ are the preferred orientations of the *i*^*th*^ pair (relative to the task orientations in the case of the task-dependent loadings), *μ*_*k*_ is a pair of orientations defining the location of the *k*^*th*^ basis function (fit to the data), and *κ*_*k*_ is the width of the *k*^*th*^ basis function (fit to the data). Because each basis function has an independent width and location fit to the data, the model predictions are non-linear functions of the parameters, unlike in the previously described regression model. Furthermore, the fitting surface has many local minima because the basis functions can simply be permuted to produce an identical model. Therefore, the optimal parameters were identified using numerical optimization with an array of starting points to identify a globally optimal solution. Since each basis function required four parameters (amplitude, width, and location in two dimensions), the total number of parameters was 4*m+1, where m is the sum of the number of allowed fixed and task-dependent basis functions and we add an additional parameter for the model constant.

### Simple parametric models

We modeled the observed set of *r*_*sc*_ values using two simple parametric models: a “single eigenvector” model and a “diagonal ridge” model. In the “single eigenvector” model, each observation *y*_*i*_ was modeled as the outer product of an eigenvector *X*,evaluated at the relevant pair of orientations. The eigenvector was parametrized as the difference of two von Mises functions separated by 90°:

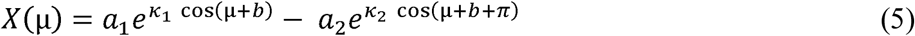

where *μ* is the difference in preferred orientation and the task orientation (in angle-doubled radians), the asare the amplitudes to be fit, the *κ’s* are the widths to be fit, and *b* is the offset of the eigenvector peak and trough from the task orientations (allowing a mismatch between the model eigenvector and the task, and also fit to the data). An observed *r*_*sc*_ value *y*_*t*_ was described as:

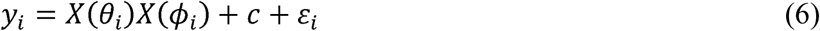

where *θ*_*i*_ and *ϕ*_*i*_ are the task-aligned preferred orientation of the pair and *c* is a model constant. The model contained six total free parameters which were fit using gradient descent to minimize the squared error in the *r*_*sc*_ predictions.

In the “diagonal ridge” model, *r*_*sc*_values were modeled as a decaying function of the difference in preferred orientation, independent of task. The dependence was modeled as a von Mises function. A given *r*_*sc*_ value *y*_*i*_ was modeled as:

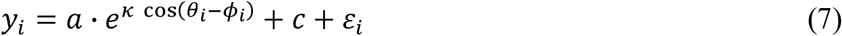

where *θ*_*i*_ and *ϕ*_*i*_ are the preferred orientation of the pair, *c* is a model constant, and *α* and *κ* parameterize the von Mises function. The model contained three total free parameters which were fit using gradient descent to minimize the squared error in the *r*_*sc*_ predictions.

### Estimating explainable variance

While the above models did not explain more than a small percentage of the variance of the raw observed *r*_*sc*_ values, this is not surprising as the raw correlation data do not vary smoothly with preferred orientation (reflecting both noise, and the fact that *r*_*sc*_ is known to depend on parameters other than orientation. ^3,22,23^). For this reason, we measured goodness-of-fit relative to an estimate of the explainable variance, which we took as the variance explained simply by a smoothed version of the raw data (sum of values in fixed and task-aligned matrices was 3.6%). Smoothing was performed with a von Mises kernel, with width chosen to maximize variance explained.

### Eye movements

Both animals tended to make anticipatory microsaccades near the end of the trial that predict their upcoming choice, consistent with a prior study^31^. This raised the possibility that choice-related eye movements gave rise to task-dependent changes in retinal input that explained the correlated fluctuations we observed. To rule this out, we measured the task-aligned *r*_*sc*_ matrix using a subset of trials on each session for which fixational eye position was not predictive of choice. To identify these trials, we used linear discriminant analysis (LDA) to predict the subject’s choices using the time series of mean binocular eye-position recorded on each trial. Then, we iteratively removed trials, starting with those furthest from the classification boundary, until classification performance no longer exceeded chance. This analysis (Supplementary Fig. 7) was restricted to the first 1.5 seconds of the trial, because we found that considering later time points (where most anticipatory microsaccades occurred) required discarding too many trials.

### Choice probability predictions

Choice Probability was calculated in the standard way^35^. We only used 0%-signal trials, as the uneven choice distributions elicited by signal trials yield noisier CP measurements. Assuming feedforward pooling with linear readout weights, the relationship between the covariance matrix for a population of neurons, the readout weight of each neuron, and the Choice Probability (*CP*) of each neuron is:

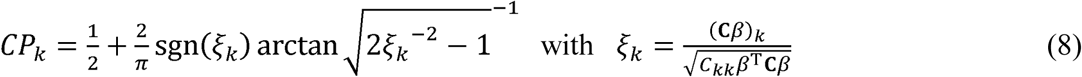

where *CP*_*k*_ is the CP of neuron *k* with respect to choice 1, *β* is the vector of readout weights and **C** is the 17 covariance matrix. We used this known relationship to quantify the CPs that would be associated with the *r*_*sc*_ structure we observed and the fixed and task-dependent components we identified, assuming only a feedforward source of CP (Fig. 8). CPs, *r*_*sc*_ structure, and readout weights were described as task-aligned functions of preferred orientation. This is equivalent to assuming a population of infinite size that is homogeneous at a given orientation. For the fixed component of *r*_*sc*_, which was indexed relative to raw orientation preferences, we generated a task-aligned version by substituting the observed *r*_*sc*_ values with model fits (using only a fixed component of the model) and then generating a smoothed task-aligned matrix as in Fig. 2e, using these substituted values. To guarantee real-valued CPs on [0,1], we performed the calculations using a symmetric positive definite approximation^55^ of the *r*_*sc*_ matrices, which introduced negligible error.

Since the readout weights were unknown, we generated a random distribution of 1000 plausible readout weight profiles that could support task performance. To generate a sample from this distribution, we started with a vector of random weights (drawn from a normal distribution) and applied the 90° symmetry inherent in the task, such that *β*_*θ*_ = −*β*_*θ*+90_, where *β*_*θ*_ is the weight assigned to neurons with task-aligned preferred orientation 9.Then, we smoothed with a wrapped Gaussian kernel with 15° s.d. and excluded profiles which did not have a circular mean within 22.5° of choice 1 (0°). In practice, we found the CP predictions to be insensitive to the readout weights (Supplementary Fig. 9), which is not surprising for a nearly rank-1 matrix (since for exactly rank-1 matrices, the CPs are independent of the weights)^17^.

We can use correlations interchangeably with covariances in Eq. 8, under the simplifying assumption that the variance is uniform as a function of preferred orientation. If **Σ** is the correlation matrix for a population with uniform variance a,then it follows that:

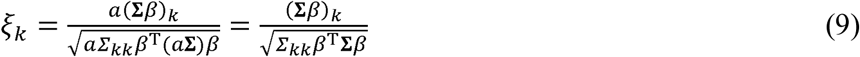

where *Σ*_*kk*_ = 1 for all *k*.We felt that spike-count variance that depended systematically on preferred orientation was unlikely to be a feature of the V1 representation, and thus that the advantages of using correlations outweighed the cost.

Noise in the decision process after pooling (pooling noise) has the effect of uniformly scaling down CPs, such that *ξ*_*K*_ in Eq. 8 is substituted with:, 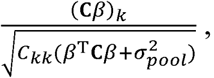, where 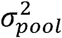 is the variance of the pooling noise^6^. We found that non-zero pooling noise was needed to avoid overestimating the magnitude of CP from the observed correlation structure. We used a fixed value of pooling noise in our predictions such that the average squared difference between the CP profile predicted from the observed correlation matrix and the observed CP profile was minimized. Empirically, we found that pooling noise variance of 0.6 was optimal. Since our spike counts were normalized to have unit variance, this implies pooling noise whose variance is 60% of the average spike-count variance of single neurons. This should be interpreted with care, as overestimation of CPs may also be an artefact related to the assumption of a homogeneous population^17^. Alternatively, the need to invoke pooling noise may be due to nonuniform sensory integration across the trial, which is distinct but which would also have an attenuating effect on CP when measured over the entire trial.

### Statistics

Statistical tests were performed non-parametrically using bootstrapping or other resampling methods, as described, with 1,000 resamples. Nonparametric statistical testing is superior when the underlying distribution of the data cannot be assumed. When p-values of p<0.001 are reported, this indicates the null hypothesis can be ruled out with the most confidence possible given the number of resamples performed. In most cases, resampling was performed from the set of recorded neuronal pairs (n=811), and always with replacement. In all figures, one asterisk indicates significant at the p<0.05 level, two indicates p<0.01, and three indicates p<0.001. When standard error bars are shown, this makes the assumption of normality in the bootstrap distribution of the test statistic. However, this assumption was not formally tested. No statistical methods were used to predetermine sample sizes but our sample sizes are similar to those of previous publications^22,23,27^.

### Data availability

All relevant data are available upon reasonable request from the authors.

### Code availability

All computer code used to generate the results are available upon request from the authors.

